# Gene expression plasticity in the antibiotic environment is predominantly adaptive in the budding yeast

**DOI:** 10.1101/2025.03.14.643277

**Authors:** Yang Qian, Zhixuan Yao, Yu Yao, Ti Qin, Piaopiao Chen, Haoxuan Liu

**Affiliations:** Center for Evolutionary & Organismal Biology and the Fourth Affiliated Hospital of Zhejiang University, Zhejiang University School of Medicine, Hangzhou 310058, China; College of Life Sciences, Zhejiang University, Hangzhou 310058, China

**Keywords:** Gene expression, phenotypic plasticity, mutation accumulation, budding yeast

## Abstract

Phenotypic plasticity plays a key role in adaptation to fluctuating environments. However, its evolutionary significance remains debated, with conflicting views on whether it is actively maintained by natural selection or a neutral consequence of molecular constraints. In this study, we investigate the evolutionary role of gene expression plasticity in yeast populations exposed to antibiotic and osmotic stress. Using mutation accumulation (MA) lines to separate the effects of natural selection from genetic drift, we compare gene expression responses (referred to as plastic change) between 22 natural strains, 21 MA lines, and their progenitor under non-stressed and stressed conditions. Our results show that, in the antibiotic environment, gene expression plasticity is selectively maintained, as indicated by its reduction in magnititude, reversal in directionality, and loss of stress-responsive pathways in MA lines. In contrast, plasticity in the osmotic stress condition appears neutral, with random variation across MA lines. This study provides direct evidence for the adaptive role of gene expression plasticity in the antibiotic environment and sheds light on its molecular mechanisms.

## Introduction

Phenotypic plasticity, the ability of a genotype to produce different phenotypes in response to environmental changes, is a fundamental feature of biological systems^1^. While it is widely recognized as an important biological property, its evolutionary origins, maintenance, and role in adaptation remain controversial. A central debate in evolutionary biology concerns whether plasticity is an adaptive mechanism actively maintained by natural selection or a byproduct of molecular constraints and regulatory networks^2–7^. Specifically, the extended evolutionary synthesis challenges the traditional modern synthesis by arguing that plasticity plays a foundational role in adaptation and that its evolutionary importance has been underappreciated^8–12^. According to the **“plasticity-first” hypothesis**^12^, when an environment shifts, the reaction norm of an organism allows it to produce a novel phenotype that may be partially or fully adapted to the new conditions. In other words, plastic phenotypic changes are often necessary for survival in novel environments, providing a crucial buffer that allows populations to persist while genetic adaptation occurs^13^. For example, plasticity facilitates adaptation by pre-aligning phenotypic traits with their optimal states, thereby serving as a stepping stone for genetic adaptation^14^. However, an alternative view posits that plastic changes are often reversed rather than reinforced during evolution, suggesting that they may not always be beneficial or necessary for adaptation^15^.

Gene expression plasticity, a key aspect of phenotypic plasticity, is particularly relevant in microbial populations, where rapid transcriptional responses to environmental stress can directly impact survival and fitness^16^. By using gene expression levels as focal traits, studying plasticity at the transcriptional level provides an effective and unbiased framework for assessing its evolutionary significance across a group of traits^15,17^. Whether gene expression plasticity is selectively maintained or merely a transient response to environmental stimuli remains uncertain, and it is also unclear whether different environments yield different outcomes.

To address this, experimental evolution provides a powerful approach by directly assessing the role of selection in shaping gene expression plasticity. Mutation accumulation (MA) experiments, in which populations undergo repeated single-cell bottlenecks to minimize the effects of natural selection, provide a rigorous approach to determine whether plasticity is selectively maintained or subject to genetic drift. If gene expression plasticity is **adaptive**, selection in natural populations would typically remove mutations that reduce plasticity, preserving an optimal level of gene expression responsiveness. However, in MA lines, where random mutations accumulate primarily due to genetic drift and selection is largely absent, deleterious mutations affecting plasticity would not be purged. As a result, plasticity should **systematically decline** in MA lines due to the accumulation of mutations that impair its regulatory mechanisms; In contrast, if gene expression plasticity is **neutral**, meaning it does not provide a selective advantage or disadvantage, its presence or absence in a given strain would be governed by stochastic processes rather than selective pressures. In this case, MA lines would show random fluctuations in plasticity, with some lines exhibiting higher or lower plasticity compared to the progenitor strain, but **without any consistent directional trend**. Thus, comparing the magnitude and direction of gene expression plasticity between the progenitor strain and MA lines provides a way to infer whether plasticity is actively maintained by selection.

In this study, we investigate the adaptive significance of gene expression plasticity using yeast MA lines exposed to antibiotic and osmotic stress. We compare gene expression responses in natural strains, where natural selection remains intact, with MA lines, where selection is largely absent. Our results reveal that gene expression plasticity in the antibiotic environment is predominantly adaptive, as indicated by its reduced magnitude in MA lines and the loss of stress-responsive pathways. Conversely, expression plasticity in the osmotic stress condition appears largely neutral, with no clear pattern of selective maintenance. These findings provide empirical evidence supporting the adaptive role of gene expression plasticity in response to antibiotics, offering new insights into how microbial populations rapidly respond to environmental stressors and how selection shapes these responses over evolutionary timescales.

## Results

### 1. Study design and transcriptome sequencing of yeast MA lines and natural strains

To investigate the selective pressures acting on expression plasticity, we utilized yeast mutation accumulation (MA) lines generated in a previous study^18^. These lines were established using BY4741 as the ancestral strain and underwent approximately 1500 generations of MA in the solid nutrient-rich (YPD) medium. To accelerate mutation accumulation, the mismatch repair gene *MSH2* was deleted before MA and reinserted afterward. Consequently, each MA lines accumulated an average of ∼900 mutations.

A total of 21 MA lines, their progenitor BY4741, and 22 natural strains^19^ were collected for analysis (**Figure 1a and Supplementary Table 1**). Given the relatively short evolutionary timeframe of the MA experiment, the genetic diversity among the MA lines averaged 1.8×10□□, significantly lower than the 3.0×10□^3^ observed in the natural strains (**Figures 1b and 1c**). Transcriptional profiles for each strain were obtained under three conditions: a non-stressful medium (YPD) and two stressed environments (YPD with 0.5 M NaCl and YPD with 1 μM clotrimazole). NaCl induces osmotic stress^20^, while clotrimazole, an antifungal drug, inhibits ergosterol synthesis in fungal cell membranes^21^. As expected, growth rates were significantly reduced in stress environments, reaching only 60%-70% of those in YPD (**Figure 1d**). For each strain, transcriptome sequencing was performed in three biological replicates, yielding an average of 7.2 million reads per sample (**Supplementary Table 2**). The high consistency among biological replicates (Pearson’s *r* = 0.98, *P* < 10□^3^□; **Supplementary Figure 1**) confirmed the robustness of the sequencing data.

**Figure 1.**
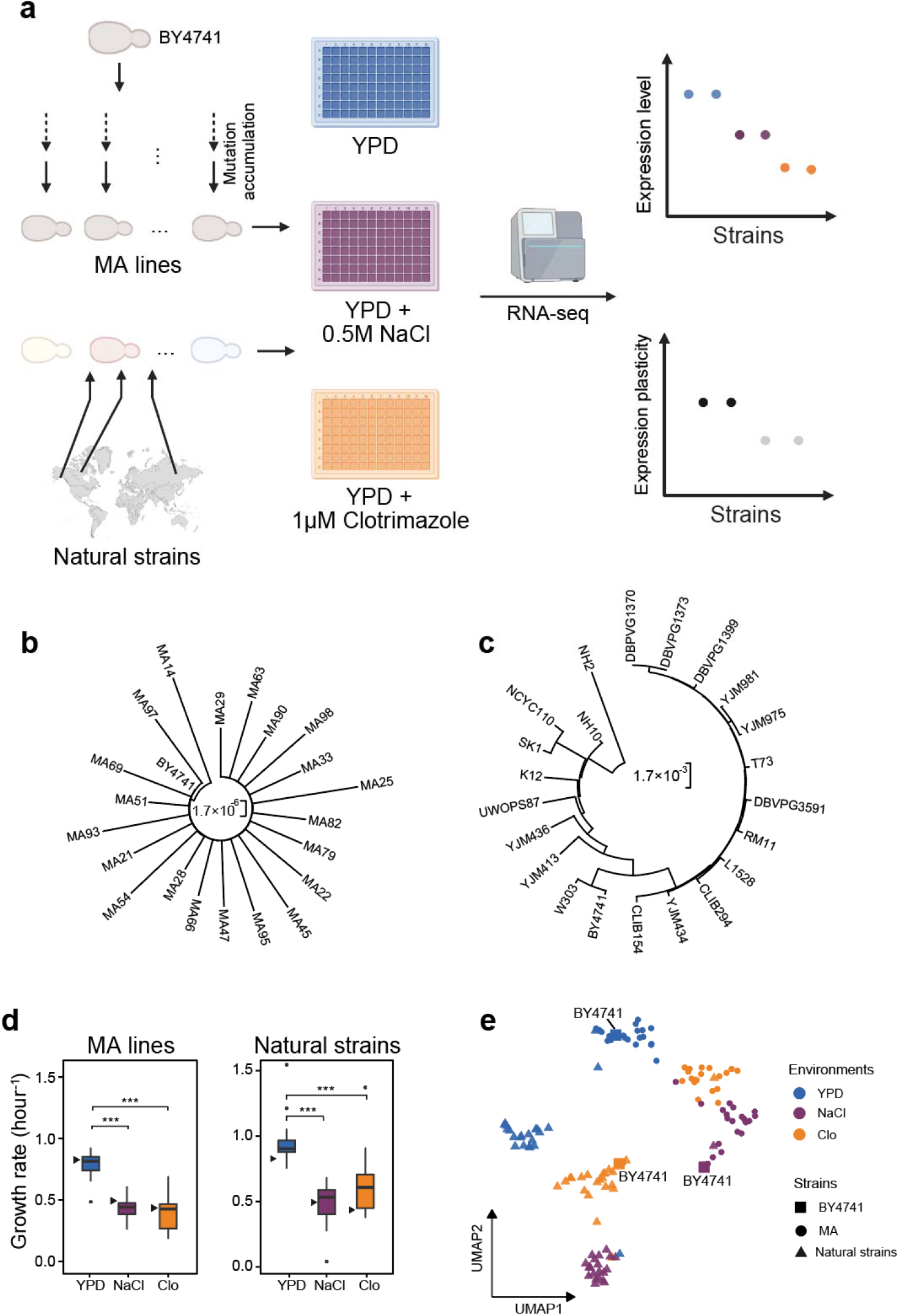
Overview of the experimental design and genetic and transcriptome relationships among the yeast strains. **a**, Schematic representation of the experimental workflow. MA strains, derived from the ancestral strain BY4741 through a laboratory mutation accumulation experiment, and natural strains, collected from natural environments, were cultured in three distinct conditions. Transcriptome were sequenced using a high-throughput platform. **b**, Phylogenetic relationship among the MA lines. **c**, Phylogenetic relationship among the natural strains. Phylogenetic trees were constructed based on all the variant sites using maximum likelihood method implemented in MEGA11. **d**, Growth rate of the MA line and natural strains in the three environments, with the ancestral strain BY4741 represented by a black triangle. **e**, UMAP visualization of gene expression profiles for the ancestor BY4741, 21 MA lines, and 22 natural strains. Colors represent different environments and shapes represent distinct strain groups.

Dimensionality reduction analysis of the transcriptome profiles revealed two major transcriptional corresponding to MA and natural strains, with further subgrouping based on environmental conditions (**Figure 1e**). These transcriptomic profiles were used to quantify gene expression plasticity by comparing expression levels across YPD and the two stress conditions (**Figure 1a**).

### 2. Reduced expression plasticity in the antibiotic environment after MA

To evaluate whether ancestral expression plasticity was selectively maintained, suppressed, or neutral, we analyzed changes in plasticity across MA lines. If ancestral plasticity was maintained by selection, MA lines would exhibit reduced plasticity due to gradual erosion of its genetic basis during mutation accumulation. Conversely, if plasticity was suppressed, MA lines would show increased plasticity, while a neutral model would predict random variation in plasticity levels relative to the progenitor.

Gene expression levels was quantified by normalizing expression levels using log_2_(FPKM+1) (**Supplementary Tables 3 and 4**)^22^. Plasticity for each gene was then calculated as the absolute difference in expression levels between stressed and YPD conditions^23^ (**Supplementary Tables 5 and 6**). To access overall plasticity, we summed the absolute plasticity across all genes for each strain. For NaCl/YPD comparisons, 9 of 21 MA lines exhibited lower plasticity compared to the progenitor, a distribution that did not significantly different from randomness (**Figure 2a**; Binomial test, *P* = 0.66). However, for Clo/YPD comparisons, 19 of 21 MA lines showed reduced plasticity (**Figure 2b**; Binomial test, *P* = 2.2×10^−4^).

**Figure 2.**
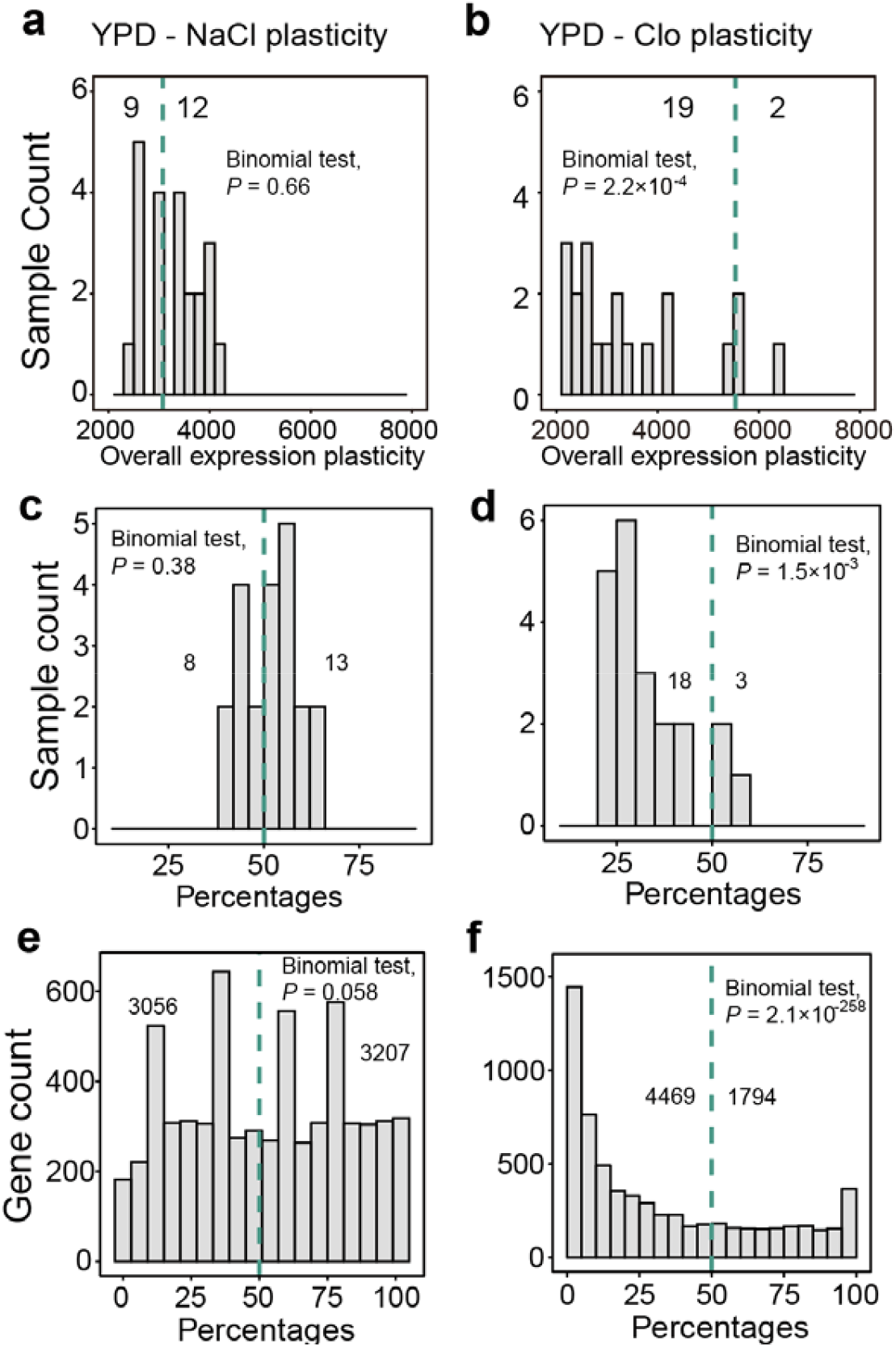
Comparisons of the level expression plasticity between MA lines and their progenitor. **a**, The overall NaCl/YPD plasticity across the MA lines. **b**, The overall Clo/YPD plasticity across the MA lines. In panels a and b, the green dashed line represents the level of plasticity in the progenitor BY4741. **c**, The percentage of genes showing elevated NaCl/YPD plasticity in comparison to the progenitor across all MA lines. **d**, The percentage of genes showing elevated Clo/YPD plasticity in comparison to the progenitor across all MA lines. **e**, The percentage of MA lines with elevated NaCl/YPD plasticity compared with the progenitor across all genes. **f**, The percentage of MA lines with elevated Clo/YPD plasticity compared with the progenitor across all genes. In panels c-f, the green dashed line represents 50%. In all panels, the numbers indicate the number of samples or genes on each side of the green line and binomial test was performed between the distribution on the two sides of the green line.

To further verify these findings, we conducted an additional analysis that weighted all genes equally. Specifically, for each MA line, we compared the expression plasticity of each gene to that of the progenitor and calculated the percentage of genes exhibiting increased plasticity. For NaCl/YPD, 8 of 21 MA lines had <50% of genes with increased plasticity, while the remaining 13 had >50%, consistent with neutrality (**Figure 2c**; *P* = 0.38). In contrast, for Clo/YPD, 18 of 21 MA lines had reduced plasticity across the majority of genes, demonstrated a reduced expression plasticity compared with their progenitor (**Figure 2d**; *P* = 1.5×10^−3^). Additionally, in 11 MA lines, fewer than 30% of genes exhibited increased plasticity, suggesting that the overall expression plasticity in the progenitor was selectively maintained.

This pattern held true when analyzing individual gene plasticity across the yeast genome. For each of the 6263 genes, we calculated the proportion of MA lines with increased plasticity compared to the progenitor. No significant bias was detected for NaCl/YPD plasticity (**Figure 2e**; *P* = 0.058). However, for Clo/YPD, a significant majority of genes in the MA lines showed reduced plasticity, providing strong evidence that plasticity was selectively maintained in the progenitor for the antibiotic environment (**Figure 2f**; Binomial test, *P* = 2.1×10^−258^).

### 3. Reduced number of significantly plastic genes in the antibiotic environment after MA

To corroborate these findings, we analyzed genes with significant plasticity (SPG) between YPD and stressed environments (FDR < 0.05). In the progenitor, 516 SPGs were identified between NaCl and YPD, with 346–1466 SPGs across MA lines (**Figure 3a**). Among the 21 MA lines, 6 exhibited fewer SPGs than the progenitor, while 15 showed more, consistent with random expectations (**Figure 3b**; *P*=0.078).

**Figure 3.**
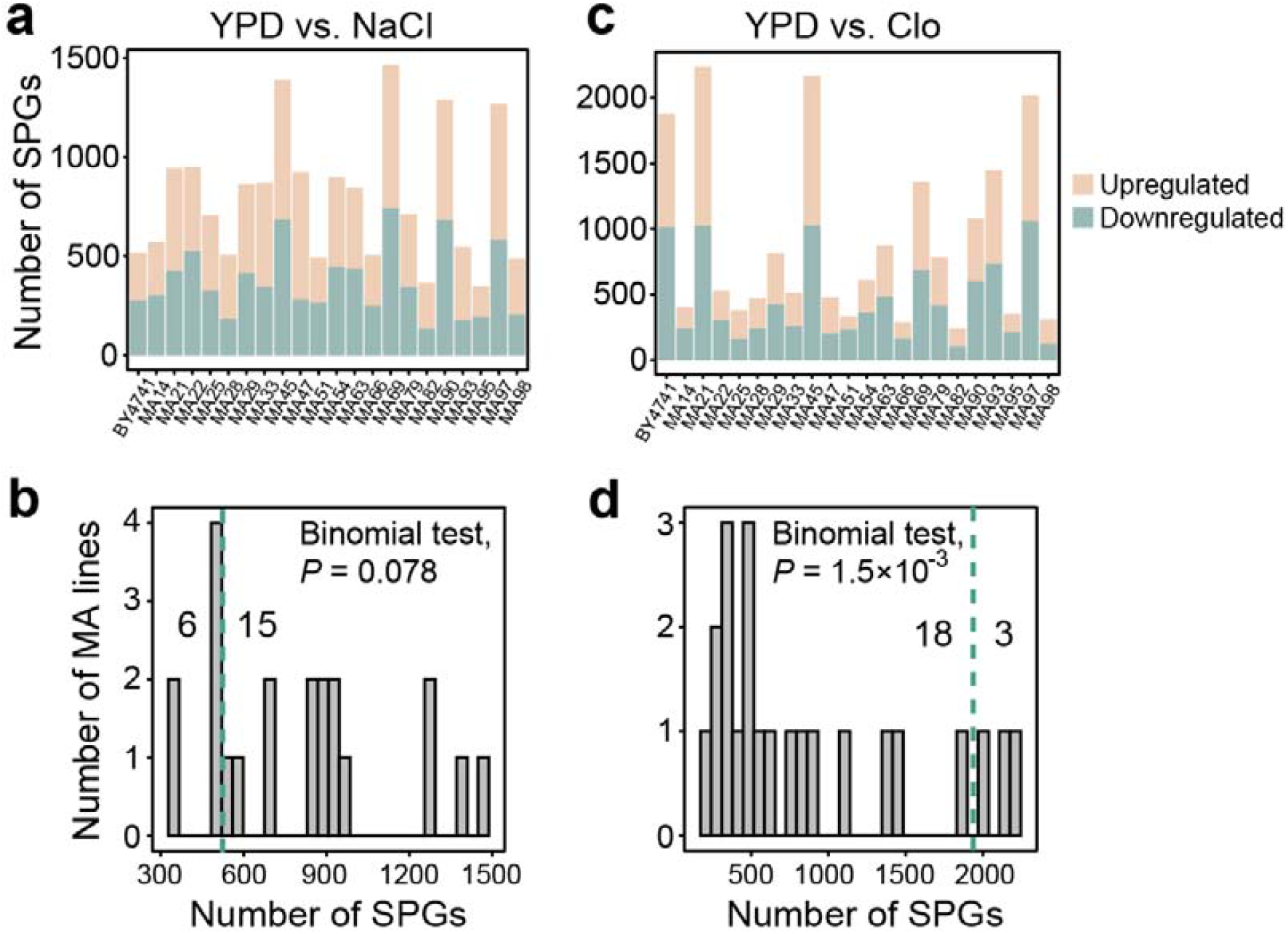
Comparison of the number of SPGs between MA lines and their progenitor. **a**, Number of SPGs between NaCl and YPD in each strain. **b**, Distribution of number SPGs between NaCl and YPD across the MA lines. **c**, Number of SPGs between clotrimazole and YPD in each strain. **d**, Distribution of number SPGs between clotrimazole and YPD across the MA lines. In panels a and c, orange represents SPGs up-regulated in the stressed environments while green represents SPGs down-regulated in the stressed environments. In panels b and d, the green dashed line represents the number of SPGs in the progenitor. The numbers indicate the number of samples on each side of the green line and binomial test was performed between the distribution on the two sides of the green line.

Between Clo and YPD conditions, 1876 SPGs were found in the progenitor, while MA lines exhibited 234–2237 SPGs (**Figure 3c**). Importantly, 18 of 21 MA lines showed reduced numbers of SPGs compared to the progenitor (**Figure 3d**; *P*=1.5×10^−3^), with an average of 840 SPGs in MA lines—less than half of the progenitor’s count. This significant reduction in transcriptional responsiveness highlights the role of selection in maintaining expression plasticity under clotrimazole stress.

### 4. Directional reversal of plasticity in the antibiotic environment after MA

While the previous analysis focused on changes in the magnitude of expression plasticity, we further investigated the evolutionary trajectory of plasticity by examining its directionality. To this end, we adapted a modified method from prior studies^6,15^ to categorize plastic expression changes into two distinct categories: **codirectional** and **reversed**. Plastic changes were classified as codirectional if the MA line retained the same directional response as the progenitor, regardless of magnitude (e.g., the expression level of a gene was higher in stressed than non-stressed conditions in both the MA line and the progenitor). Conversely, changes were classified as reversed if the direction of plasticity shifted in the opposite direction relative to the progenitor (e.g., the expression level of a gene was higher in stressed versus non-stressed conditions in the progenitor but lower under the same conditions in the MA line) (**Figure 4a**).

**Figure 4.**
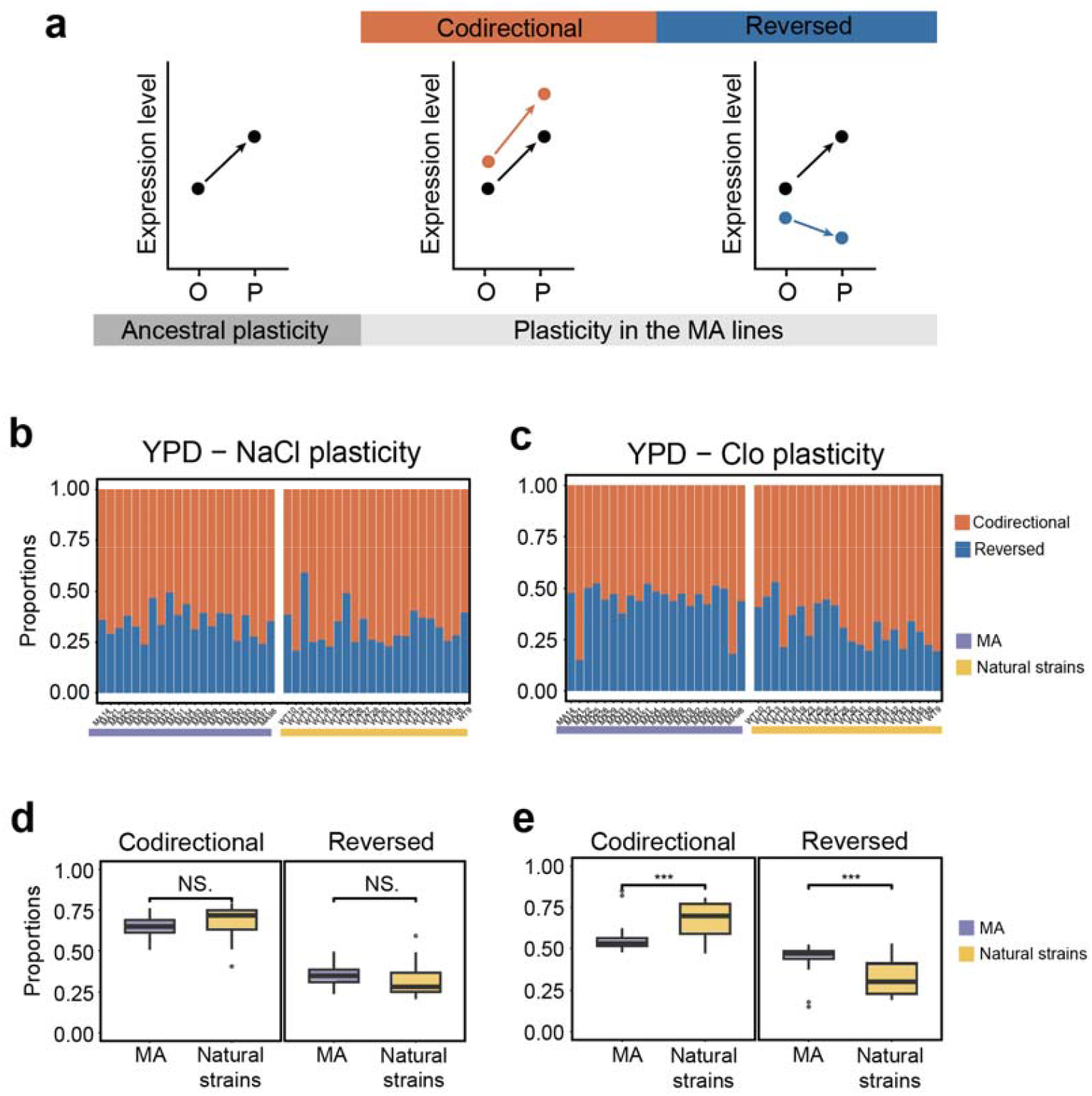
Directional change of expression plasticity in the MA lines. **a**, Classification of expression plasticity into codirectional and reversed. On the x-axis, ‘O’ represents the expression level in the original environment and ‘P’ represents the expression level in the new environment. The left subpanel illustrates the concept of ancestral plasticity and used in the right subpanel as a reference. Orange represents codirectional plasticity while blue represents reversed plasticity. **b**, The proportions of codirectional and reversed NaCl/YPD plasticity in MA lines and natural strains. **c**, The proportions of codirectional and reversed Clo/YPD plasticity in MA lines and natural strains. **d**, Comparisons of proportion of codirectional and reversed NaCl/YPD plasticity between MA lines and natural strains. **e**, Comparisons of proportion of codirectional and reversed Clo/YPD plasticity between MA lines and natural strains. In panels b-e, purple represents MA and yellow represents natural. In panels d and e, *t*-test is performed between every two bins with ‘NS.’ represents not significant and ‘***’ represents *P* < 0.001.

We hypothesized that if the directionality of ancestral plasticity was selectively maintained, reversed plasticity would occur more frequently than codirectional plasticity. This is because, in the absence of selective constraints, random mutations accumulating in MA lines are more likely to disrupt the regulatory mechanisms that sustain the ancestral plastic response. To test this, we generated a null distribution of genes between the two categories by treating the 22 natural strains as controls and comparing their plastic expression profiles with the progenitor of the MA lines. We classified all genes in the MA lines and natural strains based on plastic expression profiles in the two stressful environments (**Figures 4b** and **4c**) and compared the proportions of genes in each category between MA and natural strains.

For NaCl/YPD plasticity, no significant differences were found across the two categories between MA and natural strains (**Figure 4d**; *t*-test, all *P* > 0.2), consistent with neutral expectation. However, significant differences were observed for Clo/YPD plasticity. The MA lines exhibited significantly fewer genes with codirectional plasticity and significantly more genes with reversed plasticity compared to natural strains (**Figure 4e**; *t*-test, *P* < 0.001). The increased levels of reversed plasticity in MA lines further supports the hypothesis that expression plasticity in the progenitor was selectively maintained.

### 5. Shared pathway loss of stress response in the antibiotic environment after MA

Our results indicate that gene expression plasticity in the antibiotic environment is largely adaptive, suggesting that it provides functional benefits to yeast cells under stress. To identify the specific biological functions associated with this plasticity, we first identified genes with significant plasticity (SPGs) in the progenitor, MA lines, and natural strains. Gene Ontology (GO) enrichment analysis was then performed to identify overrepresented biological processes. If a biological process was functionally advantageous in the antibiotic environment, it would likely be present in most natural strains but lost in some MA lines. To test this, we compared the presence of progenitor-enriched biological processes between natural strains and MA lines, analyzing up-regulated and down-regulated genes separately in the antibiotic environment.

We identified 11 biological processes that were significantly enriched in the progenitor and natural strains but largely absent in MA lines (**Figure 5**; χ^2^ test, *P*-adjust < 0.05). Notably, all 11 processes were associated with genes up-regulated in the antibiotic environment, highlighting their importance under such conditions. Among the 11 enriched biological processes, they were frequently detected in natural strains (present in at least 14 of 22 strains) but rarely observed in MA lines (at most 4 of 21 lines). These processes encompassed three major functional categories: oxidative stress-related pathways, carbohydrate metabolism pathways, and purine and pyrimidine metabolism pathways. Azole antibiotics, including clotrimazole, are known to cause reactive oxygen species (ROS) accumulation in fungi^24,25^. Overexpression of oxidative stress-related pathways likely mitigates the detrimental effects of ROS accumulation. Additionally, clotrimazole inhibits glucose metabolism by targeting the key glycolytic enzyme 6-phosphofructo-1-kinase^26^. This aligns with findings that clotrimazole is more effective against fungal species like *C. albicans* that rely on glucose rather than alternative carbon sources such as lactic acid^27^. Furthermore, elevated ROS levels can lead to DNA damage^28^, necessitating enhanced nucleotide synthesis for DNA repair.

**Figure 5.**
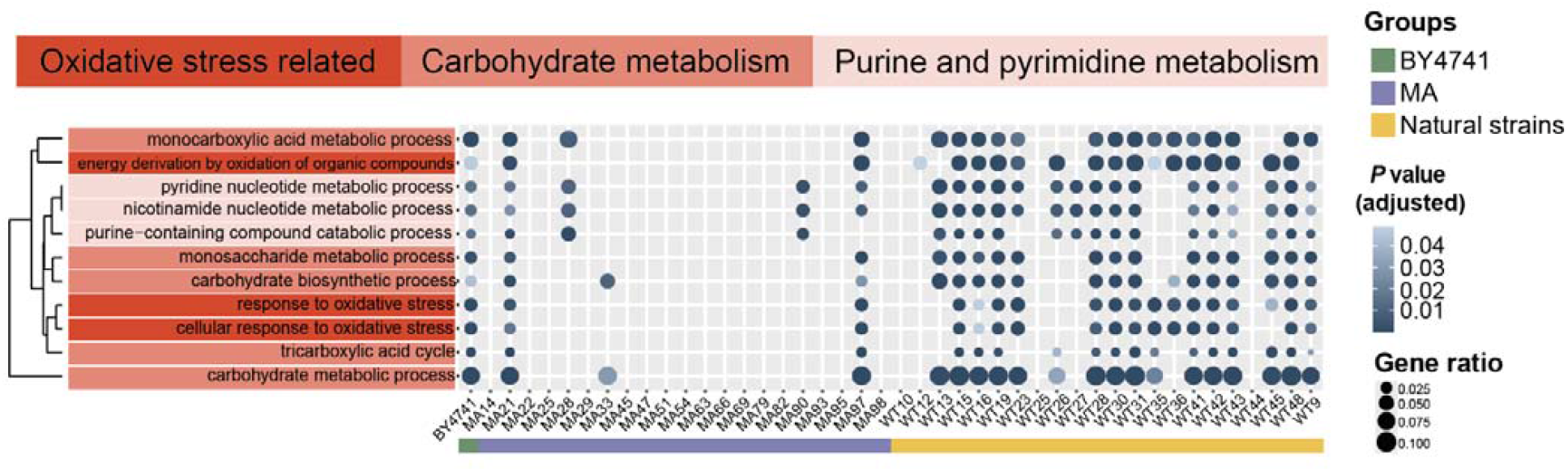
DEG enrichment in biological functions in the progenitor and natural strains but not in MA lines. Heatmap illustrating GO categories enriched in DEGs up-regulated under clotrimazole stress. Each column represents a sample and each row represents a GO category. Green, purple, and yellow represent the progenitor BY4741, MA lines, and natural strains, respectively.

In contrast to these adaptive pathways in the antibiotic environment, pathways enriched in the NaCl medium were primarily related to amino acid metabolism and showed no direct relevance to osmotic stress (**Supplementary Figure S5**). In summary, the up-regulated pathways in the progenitor and natural strains under antibiotic stress likely conferred functional advantages, while their loss in MA lines underscores the selective maintenance of these pathways in natural populations.

## Discussion

The results of this study provide compelling evidence that gene expression plasticity is not only a fundamental biological characteristic but also plays an adaptive role in response to environmental stressors such as antibiotics. Through a direct comparison between MA lines and natural strains of *Saccharomyces cerevisiae*, we demonstrated that gene expression plasticity in response to antibiotic stress is predominantly adaptive, whereas plasticity in response to osmotic stress appears neutral, showing no clear evidence of selective maintenance. These findings offer new insights into how microbial populations respond to environmental challenges and how natural selection shapes the regulation of gene expression over evolutionary time.

A key finding of this study is the significant reduction in gene expression plasticity observed in the MA lines under antibiotic stress. To quantify plasticity, we applied a widely used approach that measures differences in expression levels between stressed and non-stressed conditions^23^. To ensure the robustness of our conclusions, we also employed a normalized difference metric, where plasticity is calculated as the difference in expression levels between the two conditions divided by the sum of expression levels in both conditions. Both approaches yielded consistent results, reinforcing the conclusion that overall expression plasticity was significantly reduced in MA lines in the antibiotic environment (**Supplementary Fig. 3**). Additionally, the ancestral plasticity in the antibiotic environment was more frequently reversed (**Supplementary Fig. 4**). In contrast, under NaCl stress, no significant evidence of selective maintenance of plasticity was detected (**Supplementary Figs. 3** and **4**).

The loss of plasticity in MA lines under antibiotic stress is consistent with the idea that expression plasticity provides a selective advantage by enabling organisms to rapidly adjust their gene expression profiles in response to changing environmental conditions. This dynamic regulation likely allows yeast cells to cope with the oxidative stress and metabolic disturbances induced by clotrimazole treatment. Importantly, the specific biological pathways associated with antibiotic stress, such as oxidative stress response, carbohydrate metabolism, and nucleotide synthesis, were overexpressed in natural strains but absent or underrepresented in the MA lines, indicating the functional importance of these pathways under antibiotic pressure.

Interestingly, our analysis also revealed that plasticity in response to osmotic stress (induced by NaCl) did not exhibit the same adaptive pattern as in the antibiotic environment. The MA lines displayed a random fluctuation in plasticity, with no consistent trend towards reduced or enhanced plasticity. This suggests that gene expression plasticity in response to osmotic stress is likely neutral, as there is no clear selective pressure to maintain or modify plastic responses in the absence of natural selection. The neutral behavior of plasticity in this context contrasts with the strongly adaptive response seen under antibiotic stress, highlighting that the evolutionary significance of plasticity can vary depending on the environmental context.

Although both stressors imposed comparable selective pressures, as evidenced by similar reductions in growth rate (**Figure 1d**), their biological impact on *S. cerevisiae* may differ fundamentally. The capacity to tolerate osmotic fluctuations is likely an evolutionarily conserved trait, given the frequent exposure of yeast to NaCl in natural environments^29^. This interpretation aligns with our observation that the progenitor strain exhibited greater transcriptional robustness under osmotic stress than under antibiotic stress, as reflected by the significantly lower number of SPGs under osmotic stress (516) compared to antibiotic stress (1876). Alternatively, it is possible that yeast cells require only minimal adjustments in gene expression to cope with osmotic stress or rely on regulatory mechanisms beyond transcript abundance^30^. However, due to the limited scope and sample size of our data, these possibilities remain to be fully explored.

This study contributes to the ongoing debate in evolutionary biology regarding evolutionary origin of phenotypic plasticity. By providing direct evidence that gene expression plasticity is selectively maintained in the presence of environmental stressors, our findings support the idea that plastic expression can be selectively favored and preserved. However, the potential benefits of plasticity may arise through distinct mechanisms: by increasing cell survival rates in fluctuating environments or by aligning expression levels closer to the optimum in the new environment. Unfortunately, our data do not allow us to distinguish between these two possibilities.

## Materials and Methods

### Strains and media

This study utilized the commonly used laboratory yeast strain BY4741, 21 MA lines derived from BY4741^18^, and 22 diverse natural strains^19^ (**Supplementary Table 1**). To accumulate mutations more efficiently, the MA lines were generated in a mutator genetic background, with each line accumulating approximately 900 random mutations on average. Following mutation accumulation, the non-mutator genotype was restored for each MA line^18^. The natural strains were subjected to sporulation, polyploid selection, and genetic modifications as described by Maclean et al.^19^. Specifically, the *HO* and *URA3* genes in the natural strains were replaced with antibiotic markers, and all strains were maintained in haploid form.

Yeast strains were cultured in three media: standard YPD, YPD supplemented with 0.5 M NaCl (osmotic stress), and YPD supplemented with 1 μM clotrimazole (antibiotic stress). The YPD medium was prepared by dissolving 20 g glucose, 20 g peptone, and 10 g yeast extract in 1 L of distilled deionized water (ddH_2_O), followed by autoclaving at 121 °C for 30 minutes. For osmotic stress conditions, 29.25 g of sodium chloride was added to 1 L of YPD medium to achieve a final concentration of 0.5 M NaCl. For antibiotic stress conditions, 1.72 g of clotrimazole was dissolved in 50 mL of anhydrous ethanol, filter-sterilized, and added to 1 L of YPD medium to achieve a final concentration of 1 μM.

### Sample preparation and transcriptome sequencing

To prepare samples for transcriptome sequencing, all strains were pre-adapted in the three media (YPD, YPD + NaCl, and YPD + clotrimazole) for a full growth cycle, from initial inoculation to saturation. A 15 μL aliquot of each culture was inoculated into 985 μL of fresh medium to achieve a final volume of 1 mL. Cultures were incubated at 30 °C, and optical density (OD) was measured every one hour. Cells were harvested during the logarithmic growth phase (OD 0.4–0.6) by centrifugation at 12,000 rpm for 1 minute. The supernatant was discarded, and the cell pellets were flash-frozen in liquid nitrogen. Each strain was cultured in triplicate for each medium condition.

RNA extraction, reverse transcription, library preparation, and high-throughput sequencing were performed by Genewiz (https://climsprod.genewiz.com.cn). Sequencing was conducted on the Illumina NovaSeq 6000 platform using a PE150 configuration, generating an average of 7.2 million paired-end reads per sample (**Supplementary Table 2**).

### Data analysis

Raw sequencing data were preprocessed using fastp (v0.23.2) ^31^ to remove low-quality reads with default parameters. The Saccharomyces cerevisiae reference genome (R64) and corresponding annotation files were obtained from the Yeast Genome Database (https://www.yeastgenome.org/). Filtered reads were aligned to the reference genome using HISAT2 (v2.2.1) ^32^ and sorted using SAMtools (v1.20) ^33^. Gene-level read counts were quantified using featureCounts^34^ implemented in the Subread package (v2.0.1). Normalized expression levels, expressed as fragments per kilobase per million (FPKM), and significant plastic genes (SPGs) were defined as genes with expression level differ significantly between two environments and were identified using the DESeq2 package (v1.42.1) ^35^ in R. Gene Ontology (GO) enrichment analysis was performed using clusterProfiler (v4.10.1) ^36^.

Expression plasticity was quantified using two methods. First, FPKM values for each gene were normalized using log_2_(FPKM + 1) to represent gene expression levels (**Supplementary Tables 3 and 4**). Expression plasticity was calculated as the difference between expression levels under stress and non-stress conditions (**Supplementary Tables 5 and 6**). To ensure robustness, a second method was employed, where expression plasticity was calculated as the normalized difference, which is the difference between stress and non-stress expression levels divided by the sum of the two expression levels.

### Growth rate measurement

Growth rates of yeast strains in the three media were measured as described previously^37^. Briefly, strains were pre-adapted in the respective medium for a full growth cycle. Saturated cultures were diluted 10-fold, and 3 μL of the diluted culture was added to 197 μL of fresh medium in a 96-well plate (final volume: 200 μL per well). Plates were incubated at 30 °C in a microplate reader, and optical density at 660 nm was recorded every 20 minutes for 72 hours. Growth rates were calculated from the resulting growth curves using a previously established protocol^37^.

## Acknowledgement

Haoxuan Liu is supported by National Key Research and Development Program of China (2024YFA1802500) and National Natural Science Foundation of China (32470646). Piaopiao Chen is supported by National Key Research and Development Program of China (2024YFA1802500) and National Natural Science Foundation of China (32470650).

## Author contributions

Haoxuan Liu and Piaopiao Chen conceived the project and designed the experiment. Yang Qian, Zhixuan Yao, Yu Yao and Ti Qin carried out the experiment and analyzed the data. Haoxuan Liu, Piaopiao Chen and Yang Qian wrote the manuscript. All authors have read and approved the manuscript.

**The authors declare no conflict of interest**.

