## Supplementary Figures for "Gene expression plasticity in the antibiotic environment is predominantly adaptive in the budding yeast"

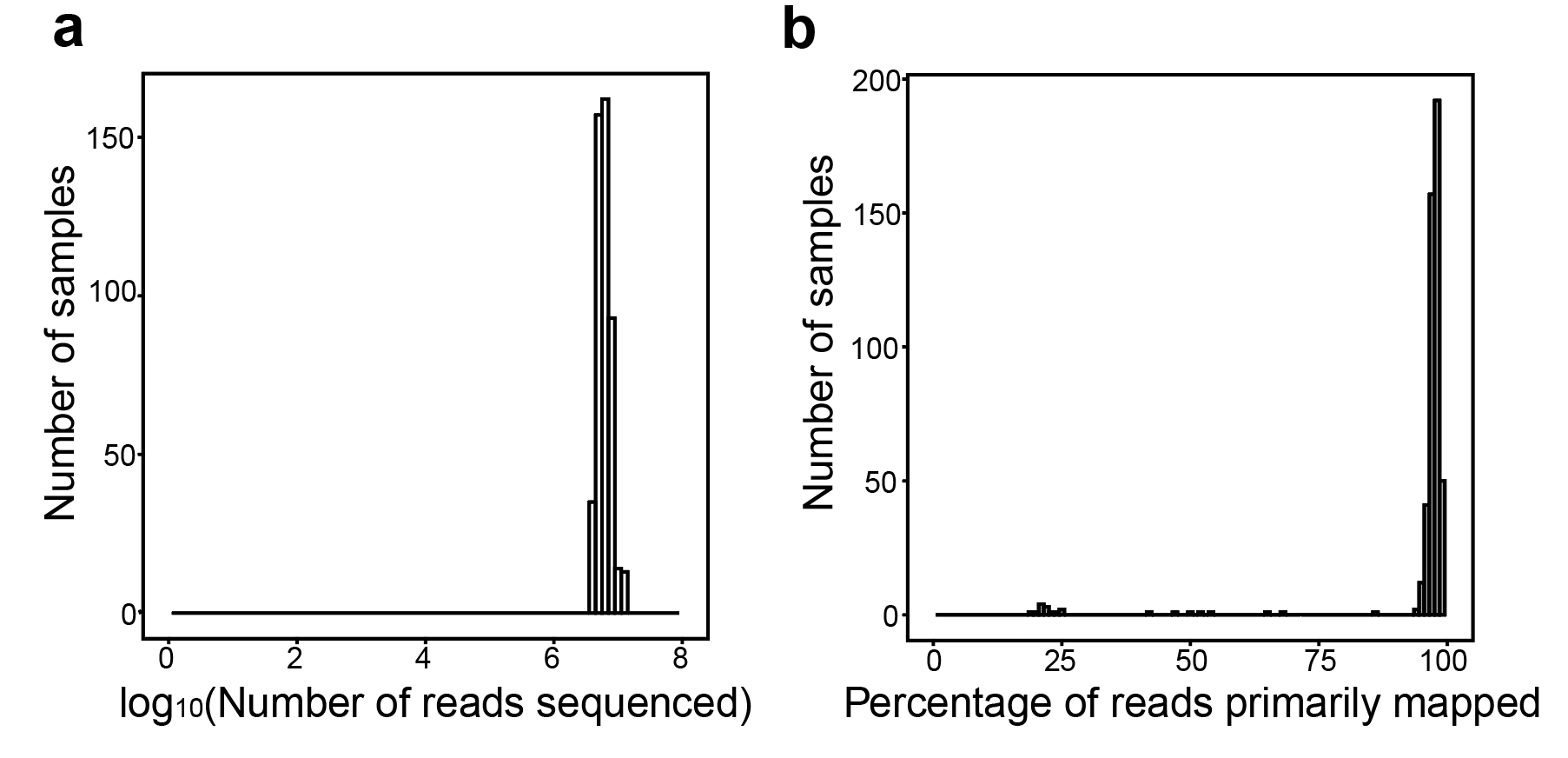

**Supplementary Fig 1. Summary statistic of the sequencing reads generated for each sample.** a, The distribution of number of pair reads sequenced per sample. b, The distribution of percentage of reads that were primarily mapped for each sample.

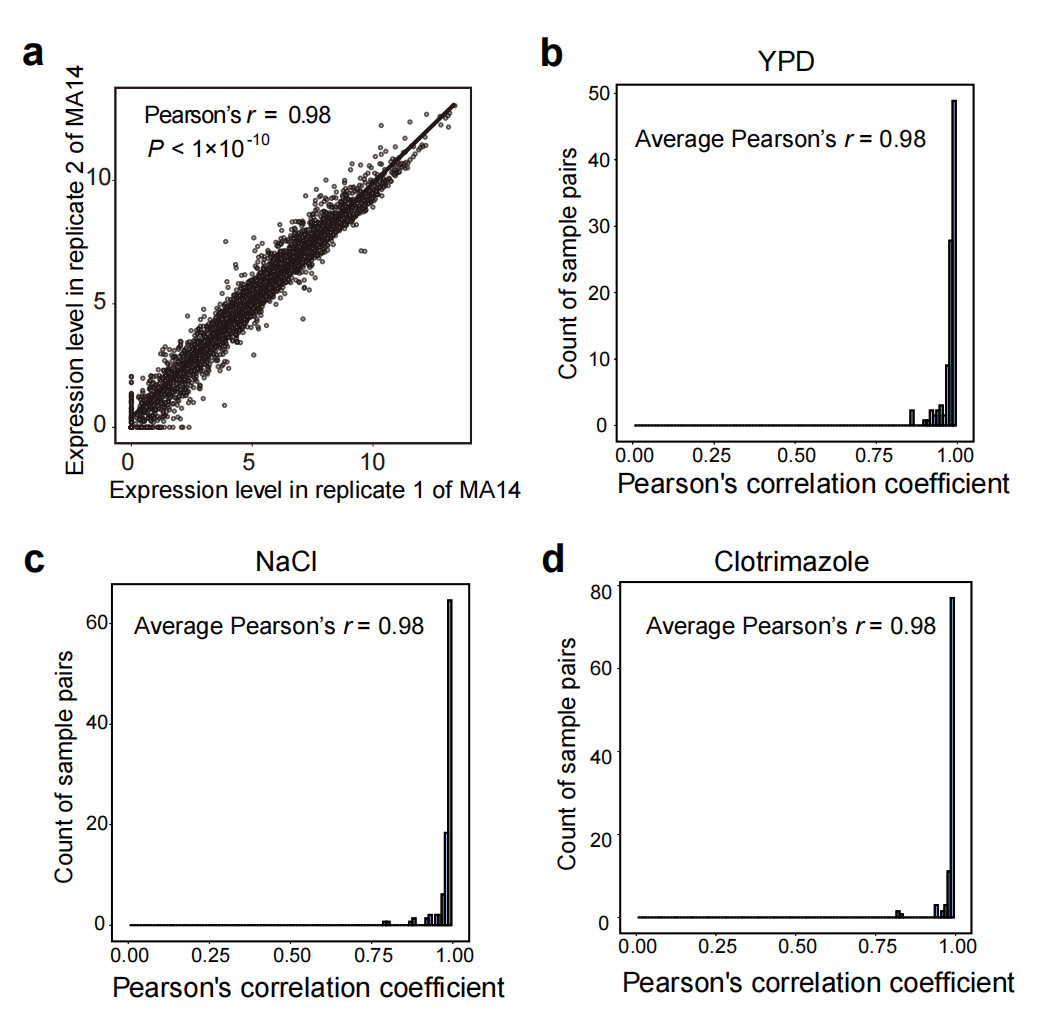

**Supplementary Fig 2. Correlation in transcriptomic profiles between biological replicates. a**, Correlation of gene expression levels between two biological replicates of MA14 across all genes. **b**, Distribution of Pearson’s r for transcriptomic profiles in YPD condition across all pair of biological replicates. **c**, Distribution of Pearson’s r for transcriptomic profiles in NaCl condition across all pair of biological replicates. **d**, Distribution for Pearson’s r of transcriptomic profiles in clotrimazole condition across all pair of biological replicates.

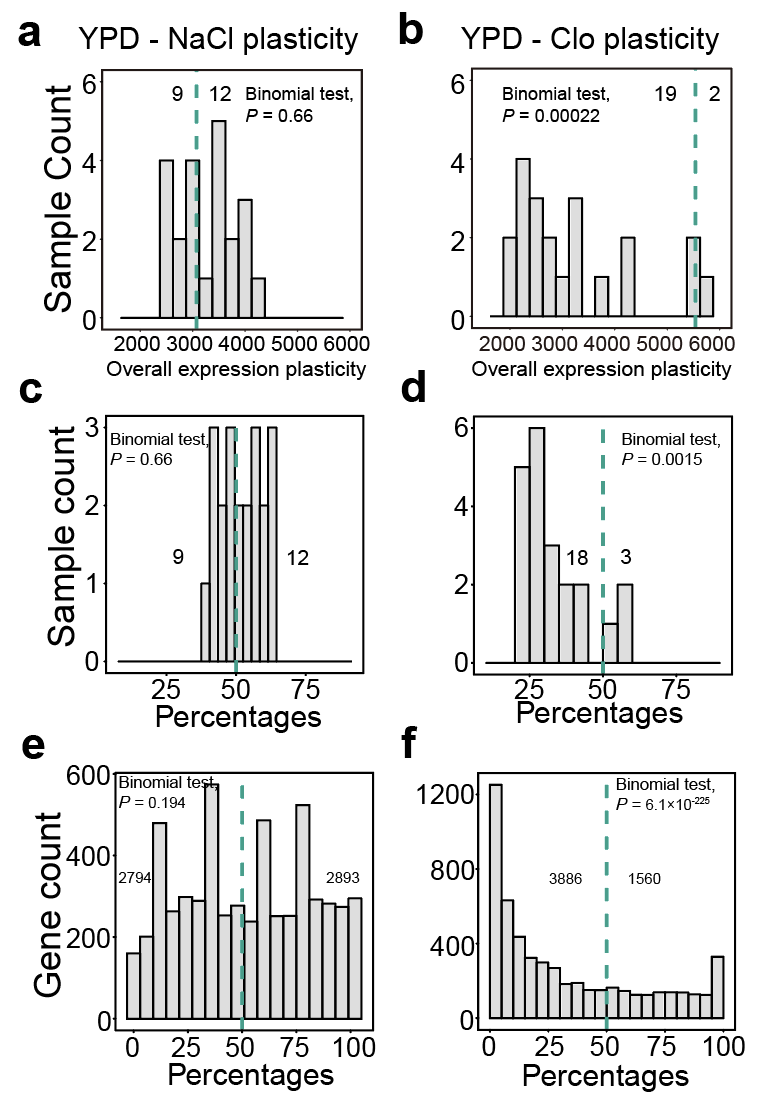

**Supplementary Fig 3. Comparisons of the level expression plasticity between MA lines and their progenitor. a**, The overall NaCl/YPD plasticity across the MA lines. **b**, The overall Clo/YPD plasticity across the MA lines. In panels a and b, the green dashed line represents the level of plasticity in the progenitor BY4741. **c**, The percentage of genes showing elevated NaCl/YPD plasticity in comparison to the progenitor across all MA lines. **d**, The percentage of genes showing elevated Clo/YPD plasticity in comparison to the progenitor across all MA lines. **e**, The percentage of MA lines with elevated NaCl/YPD plasticity compared with the progenitor across all genes. **f**, The percentage of MA lines with elevated Clo/YPD plasticity compared with the progenitor across all genes. In panels c-f, the green dashed line represents 50%. In all panels, the numbers indicate the number of samples or genes on each side of the green line and binomial test was performed between the distribution on the two sides of the green line.

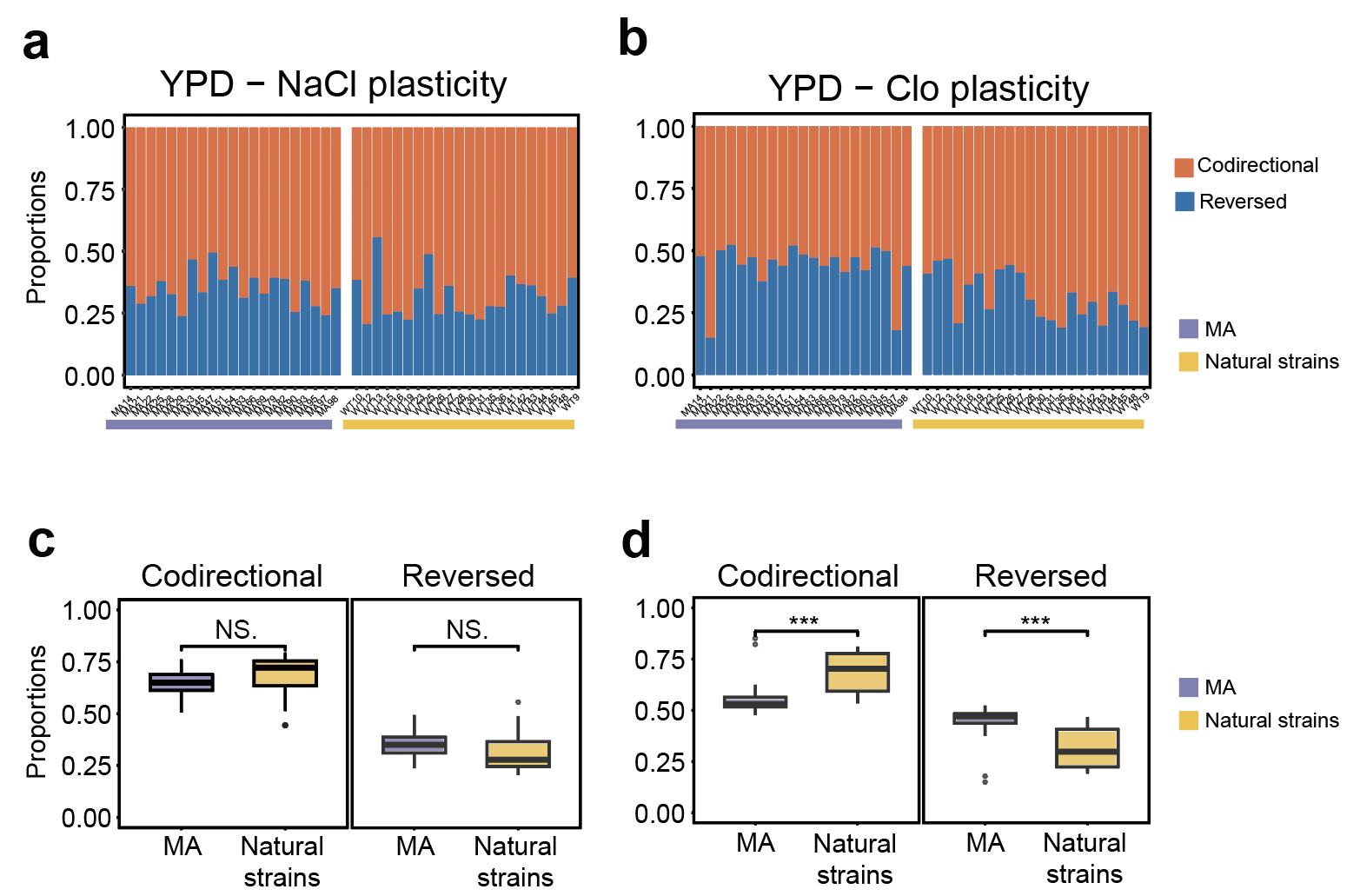

**Supplementary Fig 4. Directional change of expression plasticity in the MA lines. a**, The proportions of codirectional and reversed NaCl/YPD plasticity in MA lines and natural strains. **b**, The proportions of codirectional and reversed Clo/YPD plasticity in MA lines and natural strains. **c**, Comparisons of proportion of codirectional and reversed NaCl/YPD plasticity between MA lines and natural strains. **d**, Comparisons of proportion of codirectional and reversed Clo/YPD plasticity between MA lines and natural strains. In panels b-d, purple represents MA and yellow represents natural strains. In panels c and d, *t*-test is performed between every two bins with ‘NS.’ represents not significant and ‘***’ represents *P* < 0.001.

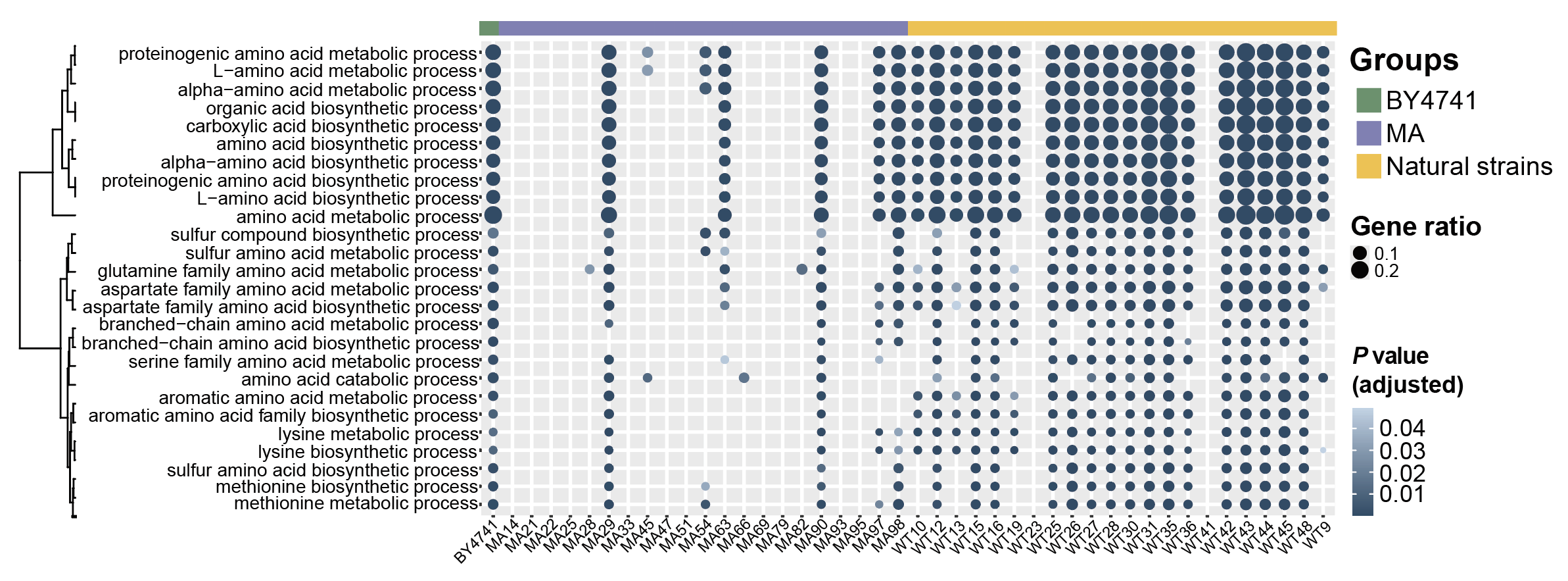

**Supplementary Fig 5. DEG enrichment in biological functions in the progenitor and natural strains but not in MA lines.** Heatmap illustrating GO categories enriched in DEGs up-regulated under NaCl stress. Each column represents a sample and each row represents a GO category. Green, purple, and yellow represent the progenitor BY4741, MA lines, and natural strains, respectively.
